# Relaxation times of ligand-receptor complex formation control T cell activation

**DOI:** 10.1101/2019.12.18.881730

**Authors:** Hamid Teimouri, Anatoly B. Kolomeisky

## Abstract

One of the most important functions of immune T cells is to recognize the presence of the pathogen-derived ligands and to quickly respond to them while at the same time not to respond to its own ligands. This is known as an absolute discrimination, and it is one of the most challenging phenomena to explain. The effectiveness of pathogen detection by T cell receptor (TCR) is limited by the chemical similarity of foreign and self-peptides and very low concentrations of foreign ligands. We propose a new mechanism of the absolute discrimination by T cells. It is suggested that the decision to activate or not to activate the immune response is controlled by the time to reach the stationary concentration of the TCR-ligand activated complex, which transfers the signal to downstream cellular biochemical networks. Our theoretical method models T-cell receptor phosphorylation events as a sequence of stochastic transitions between discrete biochemical states, and this allows us to explicitly describe the dynamical properties of the system. It is found that the proposed criterion on the relaxation times is able to explain available experimental observations. In addition, our theoretical approach explicitly analyzes the relationships between speed, sensitivity and specificity of the T cell functioning, which are the main characteristics of the process. Thus, it clarifies the molecular picture of the T cell activation processes in immune response.

## Introduction

T cells are essential components of the adaptive immune system, and they play a central role in detecting and responding to various diseases in healthy organisms [24, 32]. Activation of T cells relies on binding between a T cell receptor (TCR) and its peptide major histocompatibility complex (pMHC) on the surface of antigen-presenting cells. The inappropriate activation of T cells towards self-peptides and endogenous antigens leads to serious allergic and autoimmune responses [24]. It is known that two main factors can complicate the successful T cell functioning. First, biochemical similarity of the self-peptides and foreign peptides makes it very difficult for the T cell to elicit proper responses toward the right targets. Second, the concentrations of self-peptides in cells is known to be several orders of magnitude larger than the concentrations of foreign peptides [8, 20, 39]. The problem is that the T cells must identify very few foreign ligands in the “sea” of chemically similar self-ligands, and this should be done very quickly to avoid pathogens affecting the organisms. In other words, the successful functioning of the T cells must be simultaneously characterized by high degrees of sensitivity, specificity and speed [12].

In recent years, a substantial progress has been made in understanding the mechanisms of activation of the adaptive immune systems, and specifically for the T cells signaling [3, 6, 12–17, 21, 24, 40]. However, the molecular picture of how the absolute discrimination of self-ligands versus non-self-ligands is achieved remains not well understood. Originally, the affinity model, which assumes that the T cell activation is proportional to the probability of forming the TCR-pMHC complexes [1, 11, 34, 35, 37], has been proposed. The key prediction of this model is that the activation is a function of TCR-ligand equilibrium affinity. But weak correlations between the activation of the TCR-ligand affinity have been found in experiments [17, 34], essentially invalidating this model. The currently dominating view on the T cell activation is based on a kinetic proofreading (KPR) model [19, 24, 27]. It assumes that after the binding of TCR to pMHC several sequential phosphorylation events are taking place before the final fully phosphorylated state capable of activating the immune response is achieved. This leads to a delay between ligand binding and T cell signaling. This observation stimulated a lifetime binding concept that there is a threshold in the binding times that separates two types of ligands [17, 18, 22, 23, 30]. According to this idea, T cells do not respond to fast binders (less than ∼ 3−5 s), but they respond to ligands with longer association times. The subject remains controversial with several alternative ideas been proposed and discussed [3, 13, 15, 25]. Recent experimental studies using optogenetics techniques, however, indicated the support for the kinetic proofreading as a regulatory mechanism for the activation of the T cell receptor [38, 41], but the molecular details of the underlying processes are still not well understood.

Although the binding lifetime was found to be a good predictor for activation of T cells in some situations, in many systems it did not correlate well with the correct activation events [17, 34]. The strongest challenge to the lifetime binding concept comes from recent experimental measurements on interactions between T-cell receptors and peptide-MHC ligands [17, 34]. It was found that for fast binding peptides the lifetime failed to correctly predict the activation. In these cases, the binding lifetimes were short but the activation still took place. To explain these observations, it was argued that because of the fast on-rates there are multiple rebindings that effectively increase the overall association lifetime [17]. However, there are several issues with this approach. It assumes a slow signaling deactivation in the system which contradicts the kinetic proofreading mechanism. In addition, it was shown from experimental data that only 1 or 2 rebindings might happen, and the overall correlation between the aggregate binding lifetimes and the T cells activation improves only slightly. Furthermore, the molecular origin of the binding lifetime being the criterion for activation remains unexplained.

Here we propose a new hypothesis on the origin of absolute discrimination in T cells. We argue that the decision to activate or not to activate the immune response is governed by the characteristic time scales to form the active TCR-ligand complex that transfers the signal to the corresponding biochemical networks. The idea is developed in terms of a discrete-state stochastic model for T cell signaling that adopts a single-molecule view of the antigen discrimination process. It allows us to explicitly evaluate all relevant properties of the activation process. The hypothesis is tested then with experimental data in which the T cells response is characterized by a half maximal effective concentration (*EC*_50_), also known as a ligand potency. A clear separation of triggering the immune response for self-ligands versus foreign ligands as a function of the relaxation times is observed, supporting our theoretical predictions. The theoretical framework also provides a comprehensive description of sensitivity, selectivity and speed of T cell activation, which are the main characteristics of the antigen discrimination process.

## Model

Because of the recent experimental support for the KPR model [38, 41], in our theoretical approach we consider a discrete-state stochastic model of T cell activation as shown in Fig. 1 [27]. There are *N* states of association between TCR and pMCH that we label as states *n* for *n* = 1, 2, …, *N*. They correspond to the bound conformations with different degrees of phosphorylation. The state *n* = 0 describes the ligand and receptor being dissociated from each other. We assume that the state *n* = *N*, which is the final phosphorylated state, is the signaling state that starts the biochemical processes that lead to the T cell activation. The original TCR-pMHC complex (state *n* = 1) forms with a rate *k*_*on*_, and any of the bound conformations (states *n* = 1, 2, …, *N*) can dissociate to the state *n* = 0 with a rate *k*_*off*_ : see Fig. 1. But the rebinding of TCR and pMHC always lead to the state *n* = 1. The phosphorylation events change the state *n* to the state *n* + 1 (for *n* = 1, 2, …, *N* − 1) with a rate *k*_*p*_. This is a very simplified biochemical scheme that is believed to be capturing some relevant processes taking place during the interactions between the T cell and peptide ligands. Note that the TCR signaling transduction is a very complex biochemical process that involves multiple proteins, such as LCK, ZAP70, LAT, which recruit another effector and adaptor molecules [16].

**Figure 1.**
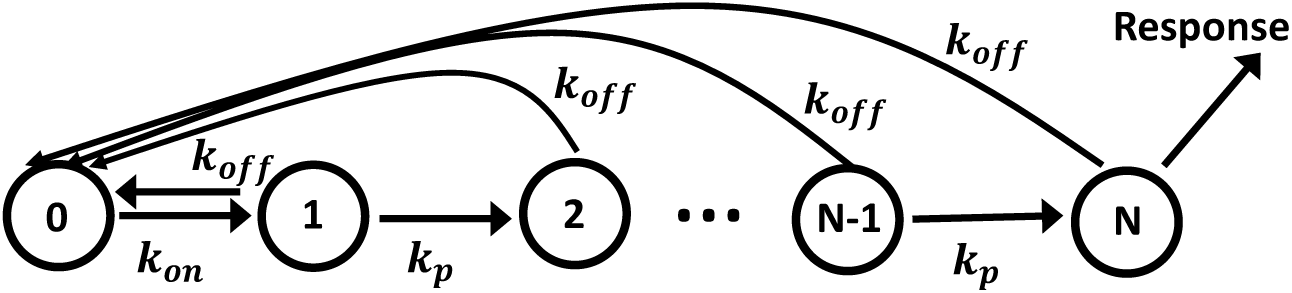
A schematic view representation of the simplest kinetic proofreading model for the antigen discrimination. Each state *n* (1 ≤ *n* ≤ *N*) corresponds to a complex between TCR and pMHC with different degree of phosphorylation. State *n* = 0 describes the unbound TCR and pMHC species. The immune response is activated when the system reaches the state *n* = *N*.

## Results

To simplify calculations, we adopt a single-molecule view of the process, i.e., the interaction between 1 TCR and 1 pMHC molecules are considered. Let us define a function *P*_*n*_ (*t*) as the probability to reach the state *n* at time *t*. At *t* = 0, the system starts in the unbounded state *n* = 0. The time evolutions of these probabilities are governed by following master equations,

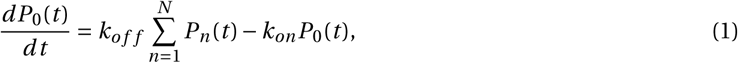

for *n* = 0, and

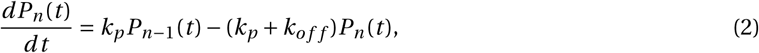

for 1 < *n* < *N* and

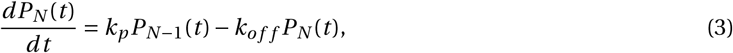

for *n* = *N*. We note that the probability functions are a subject to normalization condition, 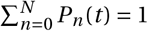. As shown in the SI, the stationary probability to reach the state *N* can be found from Eqs. (1) and (2) when the left sides of these equations are set to be equal to zero, yielding

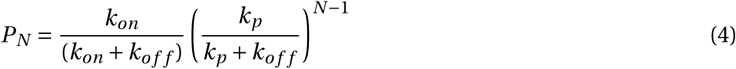

This expression has a clear physical meaning. The first term, 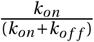, is the probability of forming the TCR-pMHC complex in any phosphorylation state, while the second term, 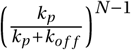, gives the probability of reaching the final activation state after *N* − 1 sequential phosphorylations [31]. Therefore, *P*_*N*_ gives the fraction of TCR-ligand fully phosphorylated complexes.

In our theoretical approach, we assume that the T cell makes the decision to activate when the system reaches the final phosphorylated state (*n* = *N* in Fig. 1). Thus, the function *P*_*N*_ can be regarded as a triggering probability such that the higher values of *P*_*N*_ correspond to the higher probability of activation. This can be tested using the experimental data as shown in Fig. 2. Using the kinetic parameters associated with different mutants of a tumor-associated protein NY-ESO-1_157−165_ interacting with 1G4 TCR [2], one can estimate *P*_*N*_ for different peptides and compare them with the half maximal effective concentrations (*EC*_50_), or ligand potency. *EC*_50_ is a concentration of the ligand that induces the activation of T cells in 50% of cases. Thus, the lower values of *EC*_50_ describe strong activation, while the higher values of *EC*_50_ correspond to weak activation response. Fig. 2 shows that the T cell activation correlates with the probability of finding the system in the final phosphorylated state *n* = *N*, and this clearly supports our theoretical arguments. Although the correlation is not perfect (*R*^2^ = 0.64), it should be noticed that experimentally measured *EC*_50_ values have large error bars (not shown in the Fig. 2) due to a variety of reasons, including fluctuations in the concentrations of participating molecules and variability in the measuring procedures.

**Figure 2.**
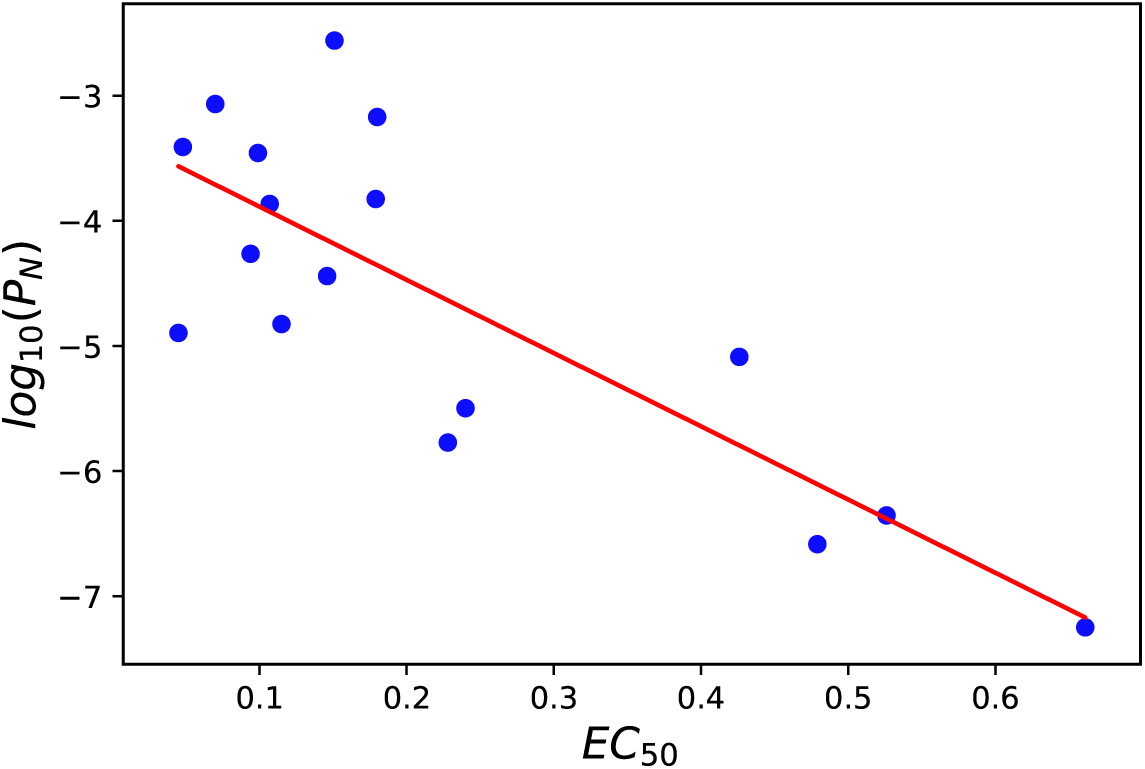
Correlations between the stationary state probability function *P*_*N*_ and the ligand potency *EC*_50_ (in units of *µ*M). Fitting analysis yields a coefficient *R*^2^ equal to 0.64. The data and kinetic parameters to evaluate *P*_*N*_ are taken from Ref [2], and they are also presented in the SI. In calculations, *N* = 6 and *k*_*p*_ = 0.1 *s*^−1^ were utilized.

Having argued that the fully phosphorylated conformation of the TCR-pMHC complex is most probably responsible for the activation of the immune response, we propose that the discrimination between self-ligands and foreign peptides is controlled by the time to reach the stationary level of this conformational state. This time, which is also known as a local relaxation time, is defined as a time to achieve the stationary probability *P*_*N*_ in the state *n* = *N* if originally the system started in the state *n* = 0 (unbound TCR and pMHC molecules). It can be explicitly evaluated using a theoretical method presented in Ref. [5]. One could define a local relaxation function for the state *n* > 0,

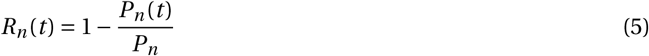

The physical meaning of this function is the relative distance to the stationary state at the state *n* at time *t*. For *n* > 0 we have *R*_*n*_ (*t* = 0) = 1, and *R*_*n*_ (*t* → ∞) = 0. It can be shown that the average time *τ*_*n*_ to reach the stationary concentration at the state *n* can be explicitly evaluated knowing the function *R*_*n*_ (*t*) (see the details of calculations in the SI). For the fully phosphorylated complex *n* = *N* we obtain,

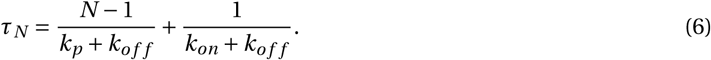

In this expression, the first term describes the time to go from the state *n* = 1 to the final state *n* = *N* via *N* − 1 phosphorylation events, while the second term is the time responsible for establishing the stationary conditions between the unbound and bound conformations in the system. Thus, our main idea is that for the ligands with the relaxation time to the final signaling state less than some threshold time *t*_0_ the activation does not happen, while for *τ*_*N*_ > *t*_0_ the immune response is activated. Experimental data also suggest that this threshold time scale is of the order of *t*_0_ ∼ 3 − 5 s [2, 17, 34].

Fig. 3 presents our theoretical predictions on the dependence of the relaxation times on the phosphorylation rate *k*_*p*_, on the complex formation rate *k*_*on*_ and on the complex dissociation rate *k*_*off*_. It shows that *τ*_*N*_ only weakly depends on the association rate, while it is much more sensitive to changes in the dissociation and phosphorylation rates. Increasing *k*_*p*_ or *k*_*off*_ lowers the relaxation time. The reason for this behavior can be understood from the chemical kinetic scheme in Fig. 1. The dominating term in the relaxation time [see Eq. (6)] is the time to move through the sequence of the phosphorylation events starting from the state *n* = 1 and finishing in the state *n* = *N*, and it depends only on *k*_*on*_ and *k*_*off*_. For larger *k*_*on*_ and *k*_*p*_, the phosphorylations are fast and this lowers the overall relaxation times, as expected. In addition, increasing *k*_*off*_ accelerates the formation of the stationary state between TCR-ligand bound and ligand unbound states.

**Figure 3.**
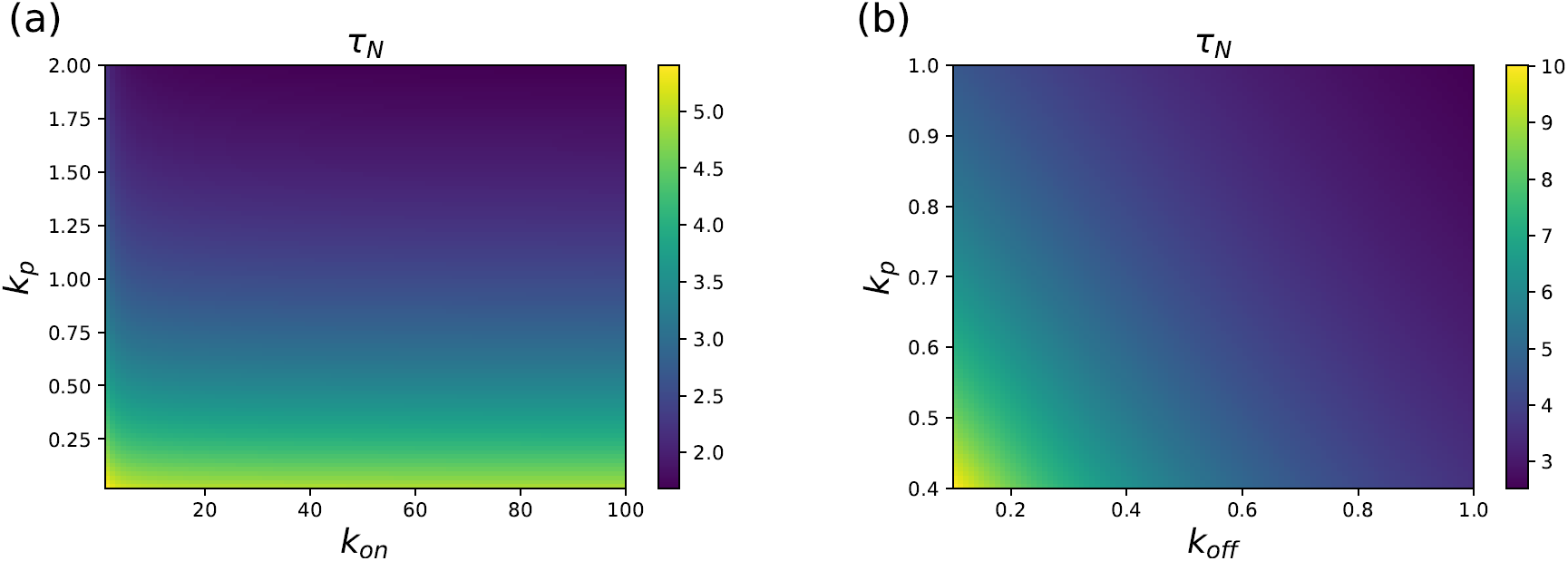
Heat maps for the relaxation times *τ*_*N*_ as a function of the transition rates in the system: (a) varying *k*_*p*_ - *k*_*on*_ (in s^−1^) parameter space (with *k*_*off*_ = 1 *s*^−1^ and *N* = 6), and (b) varying *k*_*p*_ - *k*_*off*_ (s^−1^) parameter space (with *k*_*on*_ = 100 *s*^−1^ and *N* = 6).

Now we can test our hypothesis that the amplitude of the relaxation time is the criterion for the T cell activation with experimental data. Using again the results for the protein NY-ESO-1_157−165_ interacting with 1G4 TCR [2], we present the relaxation times and the ligand potency for different T cell receptors in Fig. 4 (top). One can see that data can be clearly separated into two different groups. For *τ*_*N*_ < 5 s the value of *EC*_50_ is large, indicating that the probability of activating the immune response is low. But the situation is different for *τ*_*N*_ > 5 s: here the value of *EC*_50_ is small, and the system should be activated for these peptides. Similar observations are found for a different experimental system of *CD*4 T cells as presented in Fig. 4 (bottom). Again, the T cell activation is governed by the relaxation times. In this system, the self-ligands have *τ*_*N*_ < *t*_0_, while the foreign peptides exhibit *τ*_*N*_ > *t*_0_ with *t*_0_ ∼ 3 − 4 s. These results are generally supporting our hypothesis on the existence of the threshold time for the specific system in Fig. 4 that divides the relaxation times for the self-peptides (short times) and for the foreign ligands (long times) and determines the activation outcome.

**Figure 4.**
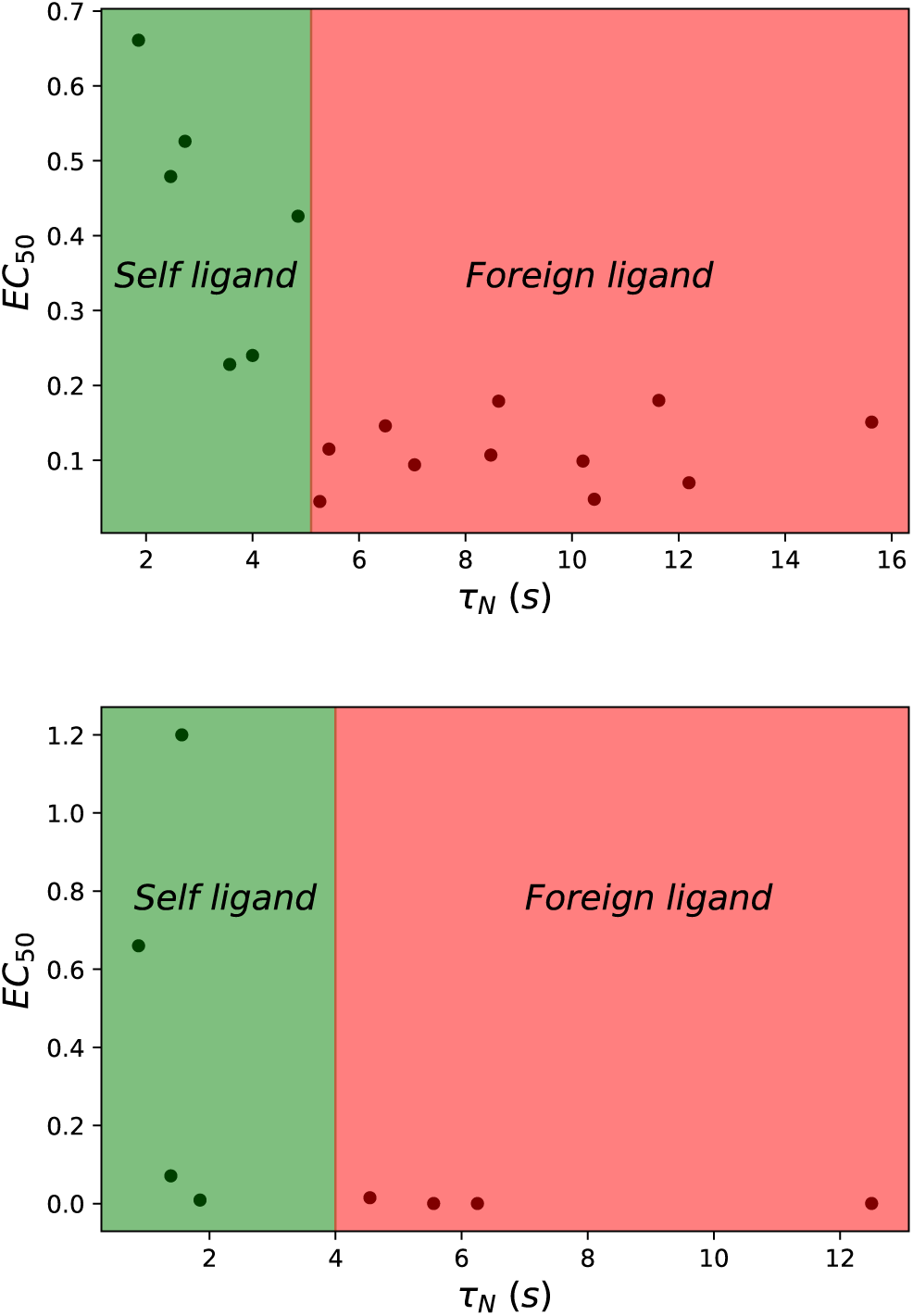
The relations between the relaxation times (*τ*_*N*_) and the ligand potency (*EC*_50_). The data and kinetic parameters to evaluate *τ*_*N*_ are taken from Ref [2] (for the top plot), and from Ref [17] (for the bottom plot). These data are also presented in the SI. In calculations, *N* = 6 and *k*_*p*_ = 0.1 *s*^−1^ were utilized.

Our theoretical approach also allows us to quantify the most important characteristics of the T cell activation process, namely, speed, sensitivity and selectivity, and the relations between them. It has been argued that the successful functioning of the immune system requires that all three properties to be in a specific range of parameters, which is called a “golden triangle” [12]. It is known that the T cells spend limited time in the vicinity of the antigen presenting cells [10, 28, 29, 33]. This requires that the T cells must act quickly to recognize and to respond to the foreign ligands. Stimulated by our theoretical hypothesis on the importance of the relaxation times, we naturally define the speed of the T cell signaling as the inverse of the time to reach the stationary concentration of the TCR-ligand activated complex,

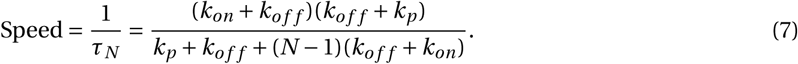

Using realistic values of the transition rates, we plot the dependence of the T cell activation speed on the number of phosphorylation steps in Fig. 5a and on the phosphorylation rate in Fig. 6a. Increasing the number of intermediate phosphorylation steps dramatically lowers the speed of activation (Fig. 5a). This is the expected result because it takes longer times for the system to reach the final signaling state *n* = *N* after the initial binding of TCR to pMHC. The trend is opposite for varying the phosphorylation rate (Fig. 6a). Increasing *k*_*p*_ accelerates the speed of the T cell activation because the system can now reach the final state *n* = *N* faster [see Eq. (6)]. However, the effect of varying the phosphorylation rate is generally weaker than changing the number of the intermediate steps in the KPR model, suggesting that for real systems the number of phosphorylation events cannot be large.

**Figure 5.**
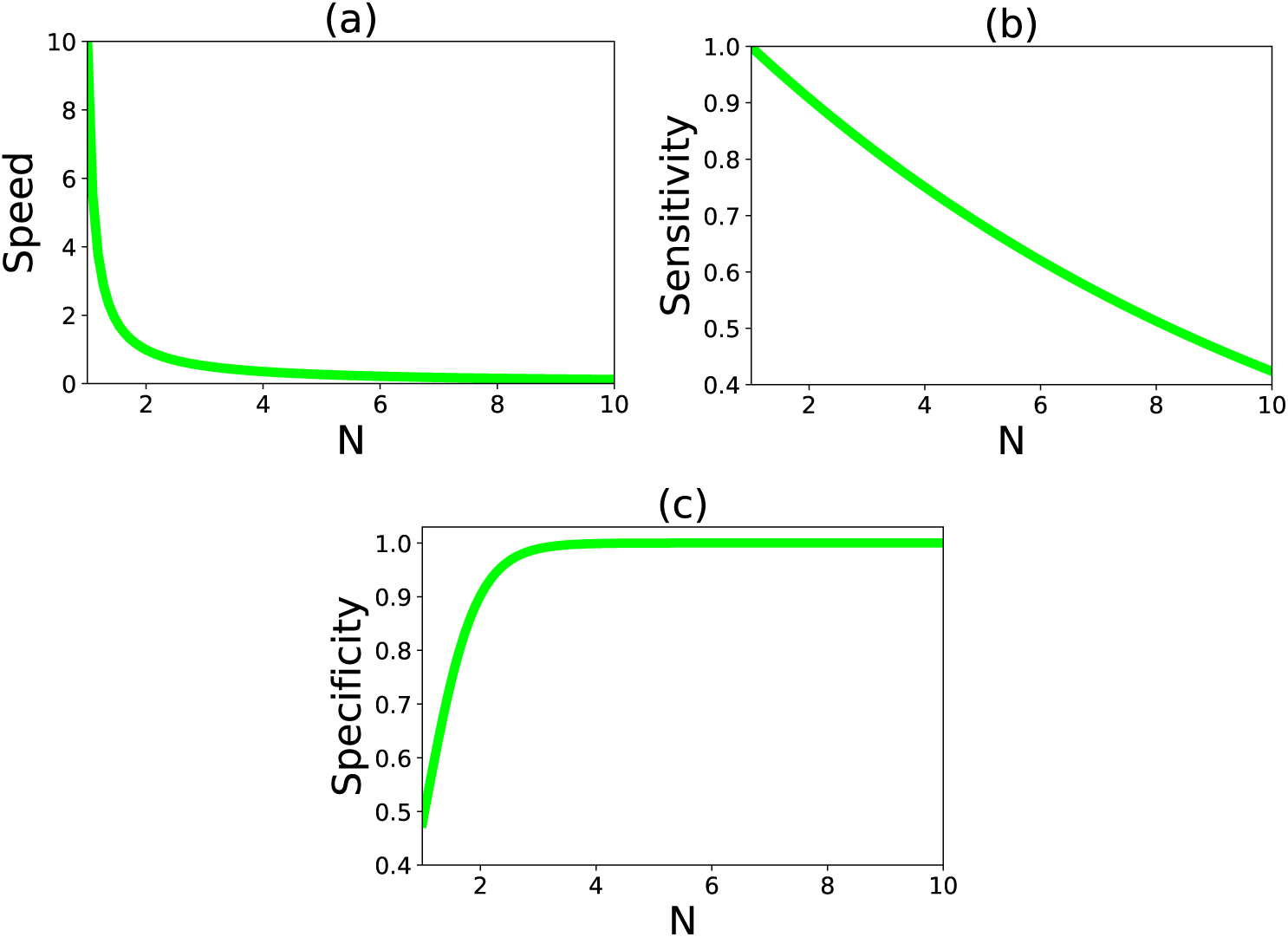
The dependence of speed, sensitivity and selectivity of T cell activation on the number of phosphorylation steps: (a) Speed for *k*_*p*_ = 1 s^−1^, *k*_*off*_ = 0.10 × *k*_*p*_, *k*_*on*_ = 10 s^−1^; (b) Sensitivity for *k*_*p*_ = 1 s^−1^, *k*_*off*_ = 0.10 × *k*_*p*_, *k*_*on*_ = 100 s^−1^; and (c) Specificity for 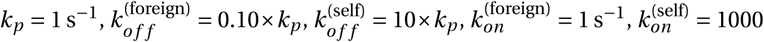 s^−1^.

**Figure 6.**
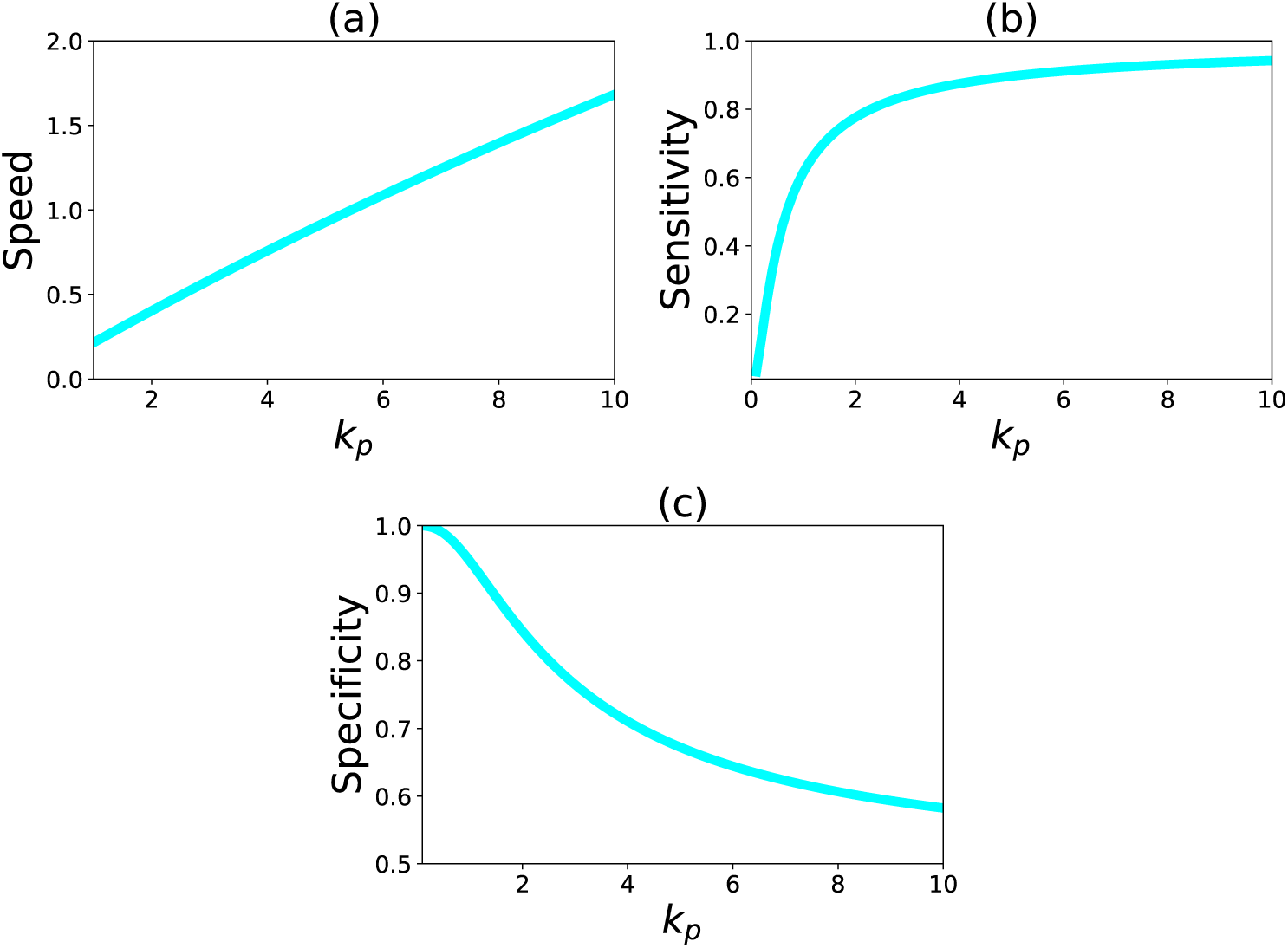
The dependence of speed, sensitivity and selectivity on the phosphorylation rate *k*_*p*_ (in s^−1^). (a) Speed for *N* = 6, *k*_*off*_ = 0.10 s^−1^, *k*_*on*_ = 10 s^−1^; (b) Sensitivity for *N* = 6, *k*_*off*_ = 0.10 s^−1^, *k*_*on*_ = 10; and (c) Specificity for 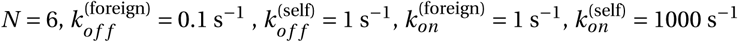.

The second major characteristics of the T cell response is sensitivity. It has been shown that the T cells recognize and respond to a very small amount of pathogen-derived ligands on antigen presenting cells [20, 36]. Sensitivity can be viewed as the probability of the achieving a state when the T cell activation starts [7]. In our formalism, this is equivalent to the probability of the stationary concentration for the fully modified TCR-ligand complex,

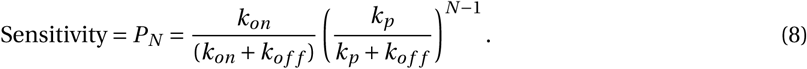

In Figs. 5b and 6b, we present the effect of varying the number of intermediate states and the phosphorylation rate on the sensitivity, respectively. Our calculations suggest that the sensitivity is lower for larger number of phosphorylation events *N* (Fig. 5b), while it strongly increases and then saturates with increasing the phosphorylation rate *k*_*p*_ (Fig. 6b). These results can be easily explained using the KPR model in Fig. 1. Increasing the number of intermediate steps lowers the probability of reaching the final state that starts the immune response. At the same time, increasing the phosphorylation rate drives the system into the direction of the final state *n* = *N*, and this should increase the sensitivity.

Specificity is the third important property of the T cell activation. Experiments suggest that even if self-ligands are presented in concentrations as high as 10^5^ times more than the foreign ligands, the T cells are able to respond only to the foreign peptides. Following the definition presented in Ref. [7], we define the specificity as the probability that the T cells produced signal is correct. Given that the TCR is exposed to specific and non-specific ligands in live cells, the specificity can be approximated as,

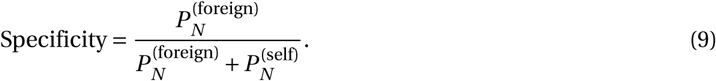

In this expression, 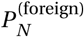 and 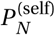 are the probabilities to reach the final signaling state for the foreign and self-ligand, respectively, for independent interactions events between TCR and pMHC molecules. In this definition, the specificity is varying between 0 (low specificity) and 1 (high specificity). Figs. 5c and 6c show how the specificity varies with the number of phosphorylation steps and the phosphorylation rate, respectively. Increasing *N* makes the T cell activation very specific. This is because at each state *n* > 0 self-ligands dissociate faster from the complex, and this lowers the probability of reaching the final signaling state *n* = *N*. The effect is stronger with larger *N*, and this improves the specificity (Fig. 5c). But increasing the phosphorylation rate lowers the specificity (Fig. 6c). In this case, both self-ligands and foreign ligands are quickly driven into the final signaling state, and there is no time for discrimination between different peptides. This result explains recent observations on tyrosine phosphorylation of the T cell adapter protein LAT at position Y132 [26]. In these experiments, the mutations that lead to faster phosphorylation kinetics have negative consequences for ligand discrimination. Our theoretical approach is able to explain from the molecular point of view.

## Discussion

In this paper, we propose a new criterion for absolute ligand discrimination in T cells activation. It argues that the relaxation time of forming the stationary concentrations of the final signaling complexes between TCR and pMHC governs the activation of the immune response. When these relaxation times are shorter than some threshold value, the signal is not activated, while for longer relaxation times the response is activated. The available experimental data generally support this picture. It should be noted that our hypothesis does not fully contradict the existing concept of binding lifetimes being the decisive factors in the activation. This is because in many situations the relaxation times and the binding lifetimes correlate with each other. But the proposed criterion works also in the situations when the binding lifetime concept fails. Thus, the hypothesis generalizes the existing views and it allows us to resolve contradictory experimental observations. It also gives a clearer molecular picture of events leading to the activation of T cells.

One of the advantages of our theoretical method is the ability to comprehensively describe the major properties of T cell activation such as speed, sensitivity and specificity. Exact analytical expressions for all these properties are presented, allowing us to analyze the intrinsic relations between them. It is shown explicitly that the specific biochemical conditions might be optimized to effectively activate the T cell response in the most efficient way. This would explain how the immune response in biological systems can be simultaneously fast, sensitive and specific. It will be also important in future to investigate the energy dissipation and other properties of the T cells activation in order to test if other features of the immune response can be optimized in the similar fashion [4, 9].

The proposed criterion that the relaxation times control the switching of the immune response agrees reasonably well with experimental observations, but it does not provide a molecular picture on the mechanisms of absolute discrimination and on the molecular origin of the threshold that separate self-ligands and foreign pathogens. The full details of downstream biochemical processes that are taking place in the immune response are still not well understood [16]. For this reason, multiple mechanisms consistent with the relaxation time criterion might be proposed. Let us suggest one such possible molecular mechanism. It is possible that *any* binding of TCR to pMHC activates some downstream biochemical cascade related to the cellular immune response. If the system reaches the final phosphorylation state (at the stationary-state level when the signal can be reliably produced) faster than the completion of the biochemical cascade activated after the binding, then the second biochemical cascade starts and it cancels the first one. As a result, the response is not activated. However, if the final signaling state reached later than the first cascade is completed, the cancellation does not happen and the immune response starts. This picture provides a possible explanation for the appearance of the threshold and its relation with the relaxation times. It will be important to get more molecular information on the processes governing the T cells activation.

Although the presented theoretical approach is able to provide a consistent description of the activation processes in T cells, it should be emphasized that the model is very simplified with multiple assumptions that need to be fully explored, and it also neglects some important features of the process. For example, it is realistic to expect a more complex biochemical network description of the interactions between TCR and pMHC [3, 16]. More specifically, the phosphorylation rate and dissociation rates might be conformational state dependent and the system might have multiple feedback loops. In the SI, the calculations for the more general situation is presented. Furthermore, our method completely neglects the role of the cellular membranes where TCR are located, cell-cell communications and cell topography during the interactions. These aspects might be important for the activation of immune response. However, despite these issues, our method provides a simple quantitative theoretical description of TCR triggering, which is based on fundamental physical-chemical principles. Another advantage of our approach is that it gives experimentally testable predictions on major properties of the T cells activation. This should clarify the molecular picture of the immune response in biological systems.

## Acknowledgement

The work was supported by the Welch Foundation (C-1559), by the NSF (CHE-1664218), and by the Center for Theoretical Biological Physics sponsored by the NSF (PHY-1427654).

In this supporting information we provide details of calculations for the equations in the main text.

## 1. Calculation of local relaxation times

We define a function *P*_*n*_(*t*) as the probability to reach the state *n* at time *t*. The dynamics in the system can be described by a set of master equations:

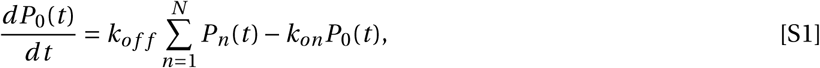

for *n* = 0, and

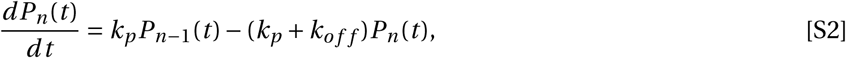

for 1 < *n* < *N* and

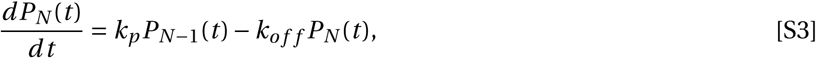

for *n* = *N*. We also have the normalization condition,

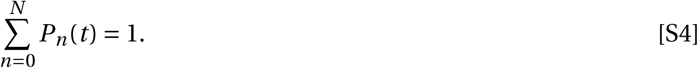

In the Laplace language, these equations can be rewritten as

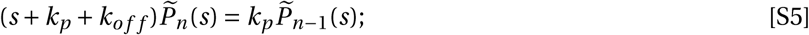

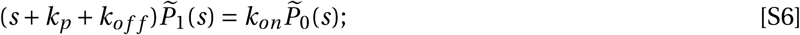

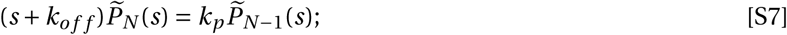

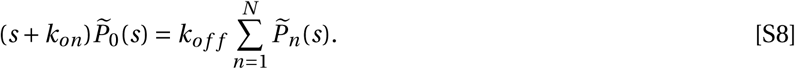

The normalization equation gives

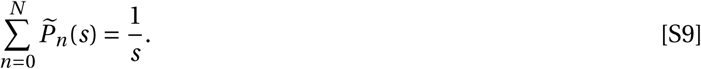

Eqs. S5, S6, S7, S8 can be solved, yielding

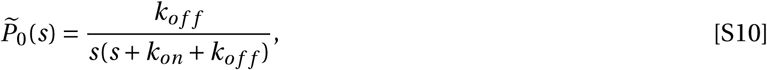

for *n* = 0; and

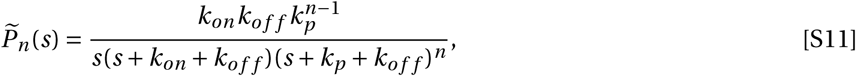

for 0 < *n* < *N*; and

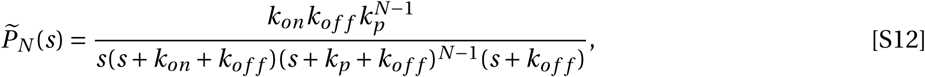

and for *n* = *N*. The stationary probabilities can be found from Eqns. S1, S2, S3 for large times when the left sides of these equations are equal to zero. We obtain then,

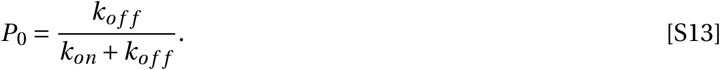

**Fig. 1.**
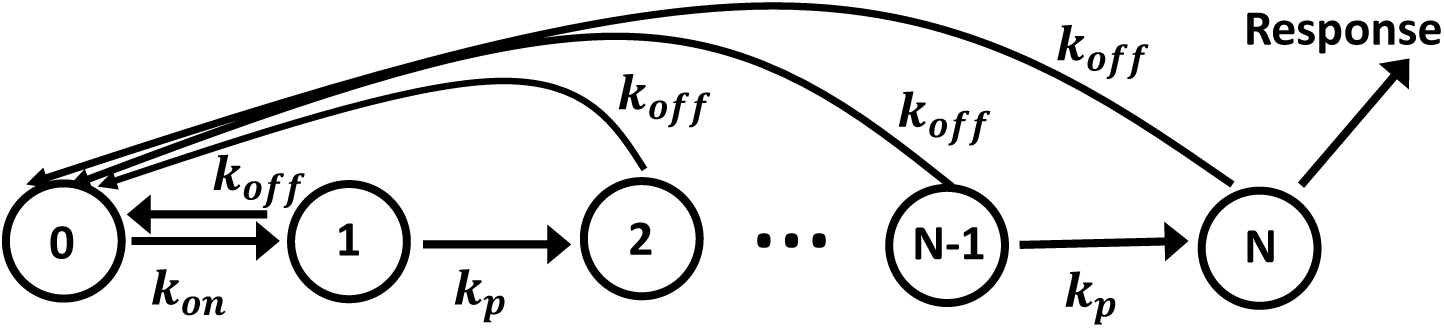
A schematic view representation of the simplest kinetic proofreading model for the antigen discrimination. Each state *n* (1 ≤ *n* ≤ *N*) corresponds to a complex between TCR and pMHC with different degree of phosphorylation. State *n* = 0 describes the unbound TCR and pMHC species. The immune response is activated when the system reaches the state *n* = *N*.

For 0 < *n* < *N* it gives

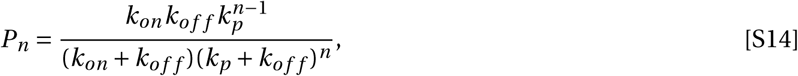

and for *n* = *N*,

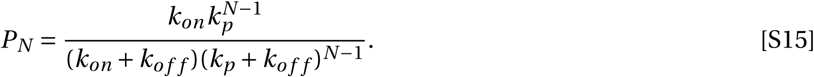

The dependence of the stationary probabilities *P*_*N*_ on various kinetic rates in the system is presented in Fig. 2. Now let us derive the times to reach the stationary states at the site *n*. We define a relaxation function *R*_*n*_ (*t*), which is given by

**Fig. 2.**
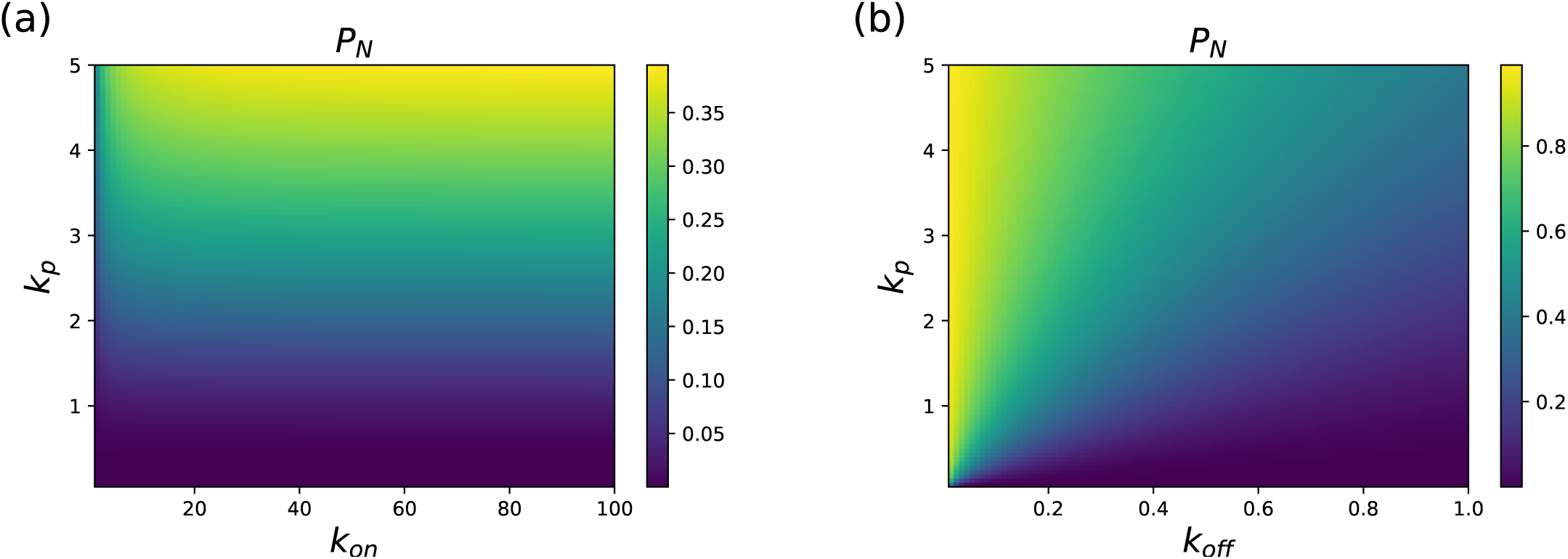
Heat maps for the stationary state probability *P*_*N*_ as a function of the transition rates in the system: (a) varying *kp* - *kon* (in s^−1^) parameter space (with *k*_*o f f*_ = 1 *s*^−1^ and *N* = 6), and (b) varying *kp* - *k*_*o f f*_ (in s^−1^) parameter space (with *kon* = 100 *s*^−1^ and *N* = 6).

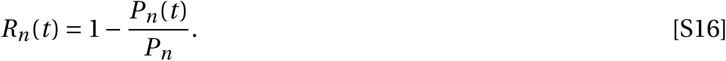

The physical meaning of this function is the relative distance to the stationary state at the state *n*. For *n* > 0, we have *R*_*n*_ (*t* = 0) = 1, and *R*_*n*_ (*t* → ∞) = 0. Therefore, it can be shown that the average time to reach the stationary concentration at the state *n* is equal to 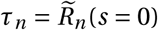. Using this expression, we obtain the times to reach the stationary states at the fully modified complex *n* = *N*,

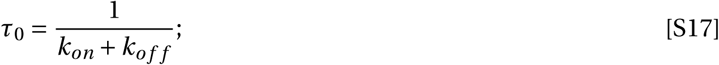

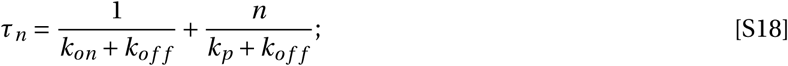

and

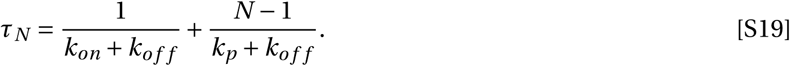

**Fig. 3.**
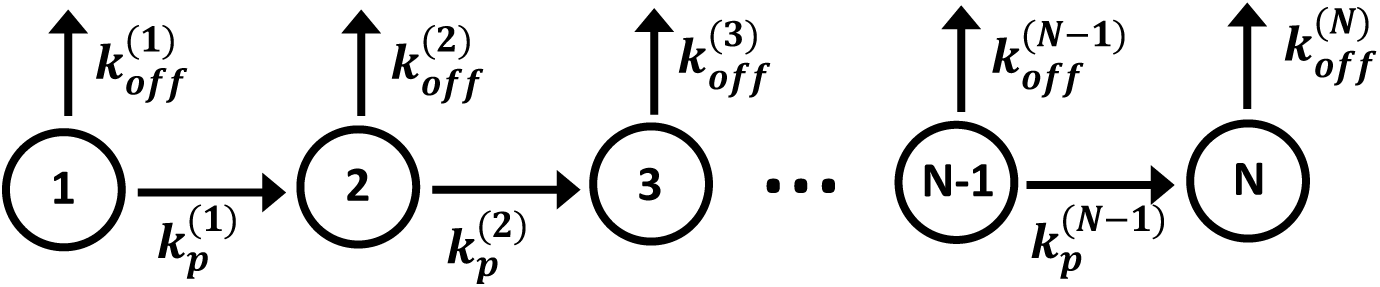
Schematic diagram for calculation of mean-first passage time.

## 2. Calculation of mean first passage times

In this section, we calculate the mean first passage time to reach a specific state. Since we only consider the first-passage times, the system dynamics become independent of the initial equilibrium binding as shown in Fig 2. Here we present a general model with inhomogeneous kinetic rates. We define *F*_*n*_ (*t*) as the probability to reach state *N* at time *t* if at *t* = 0 the system starts in the state *n* = 1. Time evolution of this function is governed by following backward master equation:

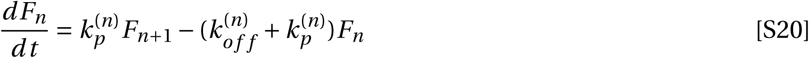

with initial condition *F*_*n*_ (*t*) = *δ*(*t*). After performing Laplace transform we obtain

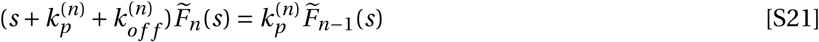

This equation leads to a full exact solution,

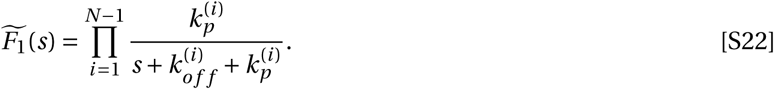

We define *T*_*n*_ as a mean-first passage time to reach the state *N* from the the state *n*. Using the probability density function *F*_*n*_ (*t*), it can be written as

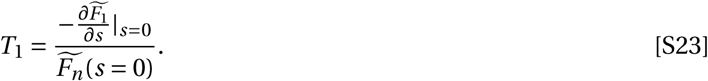

Thus, the first-passage time is given by

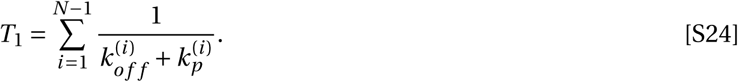

For the homogeneous kinetic rates, we obtain,

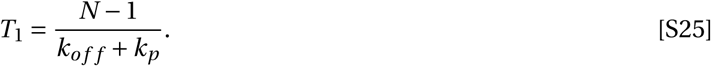

To test our theoretical calculations in the main text, we utilized the kinetic rates obtained from experimental observations in Refs. (1, 2). They are presented in Table S1 and Table S2. These data are also employed for calculating the relations between sensitivity and speed shown in Fig. S4.

**Fig. 4.**
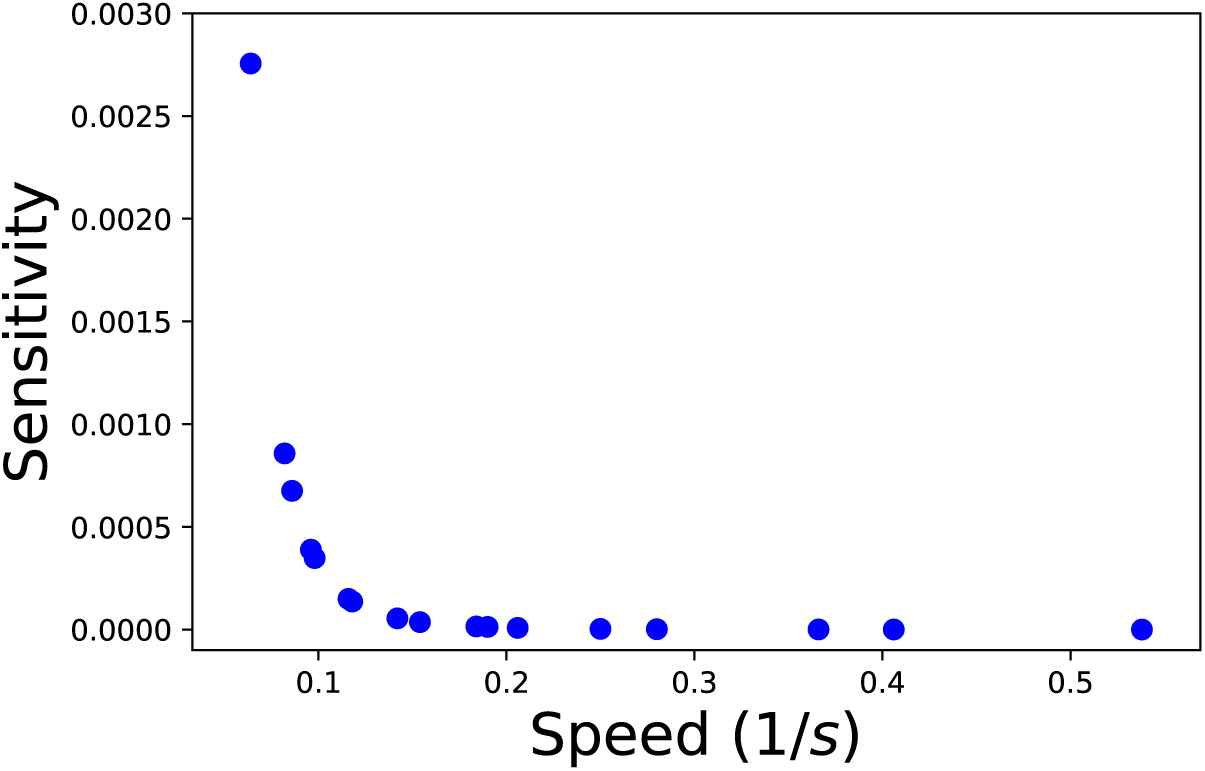
The relation between the sensitivity and speed. The data and kinetic parameters are taken from Ref (1).

**Table 1.** The data and kinetic parameters are taken from Ref (1).

| TCR | $IA^b + 3K$<br>mutation | $K_D$<br>($\mu M$) | $k_{on}$<br>(1/M.s) | $k_{off}$<br>(1/s) | $t_{1/2}$<br>(s) | Proliferation<br>$EC_{50}$ (nM) | TNF-α<br>$EC_{50}$ (nM) |
| --- | --- | --- | --- | --- | --- | --- | --- |
| B3K506 | WT | 7 | 101918 | 0.7 | 0.9 | 0.2 | 3.1 |
| B3K506 | P5R | 11 | 74654 | 0.8 | 0.9 | 0.2 | 6.0 |
| B3K506 | P8R | 13 | 64318 | 0.8 | 0.8 | 0.3 | 7.0 |
| B3K506 | P-1A | 26 | 101731 | 2.6 | 0.3 | 9.0 | 68.0 |
| B3K506 | P8A | 92 | 33370 | 3.1 | 0.2 | 1200.0 | 2210.0 |
| B3K506 | P-1K | 101 | 55149 | 5.6 | 0.1 | 660.0 | 5500.0 |
| B3K508 | WT | 29 | 10887 | 0.3 | 2.2 | 0.4 | 6.0 |
| B3K508 | P5R | 93 | 11048 | 1.0 | 0.7 | 15.0 | 87.0 |
| B3K508 | P2A | 175 | 19914 | 3.5 | 0.2 | 71.0 | 530.0 |

**Table 2.** The data and kinetic parameters are taken from Ref (2).

| peptide name | $k_{off}$ (1/s) | $EC_{50}(IFN-\gamma)$ | $k_{on}$ (1/M.s) |
| --- | --- | --- | --- |
| ESO-9C | 0.82 | 115 | 57000 |
| ESO-9L | 0.93 | 426 | 17000 |
| ESO-9V | 0.33 | 180 | 45000 |
| ESO-3A | 0.31 | 70 | 47000 |
| ESO-3I | 0.61 | 94 | 35000 |
| ESO-3M | 0.38 | 48 | 42000 |
| ESO-3Y | 1.15 | 240 | 38000 |
| ESO-4D | 2.59 | 661 | 10000 |
| ESO-6V | 0.85 | 45 | 49000 |
| ESO-6T | 1.30 | 228 | 13000 |
| ESO-7H | 1.73 | 526 | 17000 |
| A2-R65 | 1.93 | 479 | 17000 |
| A2-H70 | 0.22 | 151 | 2700 |
| A2-H74 | 0.49 | 107 | 19000 |
| A2-R75 | 0.39 | 99 | 23000 |
| A2-V76 | 0.67 | 146 | 31000 |
| A2-K146 | 0.48 | 179 | 24000 |

## References

1. S.M. M. Alam, P. J. Travers, J. L. Wung, W. Nasholds, S. Redpath, S. C. Jameson, and N. R. Gascoigne. T-cell-receptor affinity and thymocyte positive selection. Nature, 381(6583):616, 1996.

2. M. Aleksic, O. Dushek, H. Zhang, E. Shenderov, J.-L. Chen, V. Cerundolo, D. Coombs, and P. A. van der Merwe. Dependence of t cell antigen recognition on t cell receptor-peptide mhc confinement time. Immunity, 32(2):163–174, 2010.

3. G. Altan-Bonnet and R. N. Germain. Modeling t cell antigen discrimination based on feedback control of digital erk responses. PLoS biology, 3(11):e356, 2005.

4. K. Banerjee, A. B. Kolomeisky, and O. A. Igoshin. Elucidating interplay of speed and accuracy in biological error correction. Proceedings of the National Academy of Sciences, 114(20):5183–5188, 2017.

5. A. M. Berezhkovskii, C. Sample, and S. Y. Shvartsman. How long does it take to establish a morphogen gradient? Biophysical journal, 99(8):L59–L61, 2010.

6. A. K. Chakraborty and A. Weiss. Insights into the initiation of tcr signaling. Nature immunology, 15(9):798, 2014.

7. C. Chan, A. J. George, and J. Stark. T cell sensitivity and specificity-kinetic proofreading revisited. Discrete & Continuous Dynamical Systems-B, 3(3):343–360, 2003.

8. C. J. Cohen, O. Sarig, Y. Yamano, U. Tomaru, S. Jacobson, and Y. Reiter. Direct phenotypic analysis of human mhc class i antigen presentation: visualization, quantitation, and in situ detection of human viral epitopes using peptide-specific, mhc-restricted human recombinant antibodies. The Journal of Immunology, 170(8):4349–4361, 2003.

9. W. Cui and P. Mehta. Identifying feasible operating regimes for early t-cell recognition: The speed, energy, accuracy trade-off in kinetic proofreading and adaptive sorting. PloS one, 13(8):e0202331, 2018.

10. M. L. Dustin. Stop and go traffic to tune t cell responses. Immunity, 21(3):305–314, 2004.

11. L. K. Ely, K. J. Green, T. Beddoe, C. S. Clements, J. J. Miles, S. P. Bottomley, D. Zernich, L. Kjer-Nielsen, A. W. Purcell, J. McCluskey, et al. Antagonism of antiviral and allogeneic activity of a human public ctl clonotype by a single altered peptide ligand: implications for allograft rejection. The Journal of Immunology, 174(9):5593–5601, 2005.

12. O. Feinerman, R. N. Germain, and G. Altan-Bonnet. Quantitative challenges in understanding ligand discrimination by *α β* t cells. Molecular immunology, 45(3):619, 2008.

13. R. A. Fernandes, K. A. Ganzinger, J. C. Tzou, P. Jönsson, S. F. Lee, M. Palayret, A. M. Santos, A. R. Carr, A. Ponjavic, V. T. Chang, et al. A cell topography-based mechanism for ligand discrimination by the t cell receptor. Proceedings of the National Academy of Sciences, page 201817255, 2019.

14. P. François and G. Altan-Bonnet. The case for absolute ligand discrimination: modeling information processing and decision by immune t cells. Journal of Statistical Physics, 162(5):1130–1152, 2016.

15. P. François, G. Voisinne, E. D. Siggia, G. Altan-Bonnet, and M. Vergassola. Phenotypic model for early t-cell activation displaying sensitivity, specificity, and antagonism. Proceedings of the National Academy of Sciences, 110(10):E888–E897, 2013.

16. G. Gaud, R. Lesourne, and P. E. Love. Regulatory mechanisms in t cell receptor signalling. Nature Reviews Immunology, 18(8):485–497, 2018.

17. C. C. Govern, M. K. Paczosa, A. K. Chakraborty, and E. S. Huseby. Fast on-rates allow short dwell time ligands to activate t cells. Proceedings of the National Academy of Sciences, 107(19):8724–8729, 2010.

18. A. Grakoui, S. K. Bromley, C. Sumen, M. M. Davis, A. S. Shaw, P. M. Allen, and M. L. Dustin. The immunological synapse: a molecular machine controlling t cell activation. Science, 285(5425):221–227, 1999.

19. J. J. Hopfield. Kinetic proofreading: a new mechanism for reducing errors in biosynthetic processes requiring high specificity. Proceedings of the National Academy of Sciences, 71(10):4135–4139, 1974.

20. D. J. Irvine, M. A. Purbhoo, M. Krogsgaard, and M. M. Davis. Direct observation of ligand recognition by t cells. Nature, 419(6909):845, 2002.

21. G. J. Kersh and P. M. Allen. Essential flexibility in the t-cell recognition of antigen. Nature, 380(6574):495–498, 1996.

22. G. J. Kersh, E. N. Kersh, D. H. Fremont, and P. M. Allen. High-and low-potency ligands with similar affinities for the tcr: the importance of kinetics in tcr signaling. Immunity, 9(6):817–826, 1998.

23. M. Krogsgaard, N. Prado, E. J. Adams, X.-l. He, D.-C. Chow, D. B. Wilson, K. C. Garcia, and M. M. Davis. Evidence that structural rearrangements and/or flexibility during tcr binding can contribute to t cell activation. Molecular cell, 12(6):1367–1378, 2003.

24. M. Lever, P. K. Maini, P. A. Van Der Merwe, and O. Dushek. Phenotypic models of t cell activation. Nature Reviews Immunology, 14(9):619, 2014.

25. L. Limozin, M. Bridge, P. Bongrand, O. Dushek, P. A. van der Merwe, and P. Robert. Tcr–pmhc kinetics under force in a cell-free system show no intrinsic catch bond, but a minimal encounter duration before binding. Proceedings of the National Academy of Sciences, 116(34):16943–16948, 2019.

26. W.-L. Lo, N. H. Shah, S. A. Rubin, W. Zhang, V. Horkova, I. R. Fallahee, O. Stepanek, L. I. Zon, J. Kuriyan, and A. Weiss. Slow phosphorylation of a tyrosine residue in lat optimizes t cell ligand discrimination. Nature immunology, 20(11):1481–1493, 2019.

27. T. W. McKeithan. Kinetic proofreading in t-cell receptor signal transduction. Proceedings of the national academy of sciences, 92(11):5042–5046, 1995.

28. T. R. Mempel, S. E. Henrickson, and U. H. Von Andrian. T-cell priming by dendritic cells in lymph nodes occurs in three distinct phases. Nature, 427(6970):154, 2004.

29. M. J. Miller, A. S. Hejazi, S. H. Wei, M. D. Cahalan, and I. Parker. T cell repertoire scanning is promoted by dynamic dendritic cell behavior and random t cell motility in the lymph node. Proceedings of the National Academy of Sciences, 101(4):998–1003, 2004.

30. S. Qi, M. Krogsgaard, M. M. Davis, and A. K. Chakraborty. Molecular flexibility can influence the stimulatory ability of receptor–ligand interactions at cell–cell junctions. Proceedings of the National Academy of Sciences, 103(12):4416–4421, 2006.

31. C. Salazar and T. Höfer. Multisite protein phosphorylation–from molecular mechanisms to kinetic models. The FEBS journal, 276(12):3177–3198, 2009.

32. J. E. Smith-Garvin, G. A. Koretzky, and M. S. Jordan. T cell activation. Annual review of immunology, 27:591–619, 2009.

33. S. Stoll, J. Delon, T. M. Brotz, and R. N. Germain. Dynamic imaging of t cell-dendritic cell interactions in lymph nodes. Science, 296(5574):1873–1876, 2002.

34. J. D. Stone, A. S. Chervin, and D. M. Kranz. T-cell receptor binding affinities and kinetics: impact on t-cell activity and specificity. Immunology, 126(2):165–176, 2009.

35. Y. Sykulev, R. J. Cohen, and H. N. Eisen. The law of mass action governs antigen-stimulated cytolytic activity of cd8+ cytotoxic t lymphocytes. Proceedings of the National Academy of Sciences, 92(26):11990–11992, 1995.

36. Y. Sykulev, M. Joo, I. Vturina, T. J. Tsomides, and H. N. Eisen. Evidence that a single peptide–mhc complex on a target cell can elicit a cytolytic t cell response. Immunity, 4(6):565–571, 1996.

37. S. Tian, R. Maile, E. J. Collins, and J. A. Frelinger. Cd8+ t cell activation is governed by tcr-peptide/mhc affinity, not dissociation rate. The Journal of Immunology, 179(5):2952–2960, 2007.

38. D. K. Tischer and O. D. Weiner. Light-based tuning of ligand half-life supports kinetic proofreading model of t cell signaling. eLife, 8:e42498, 2019.

39. J. J. Unternaehrer, A. Chow, M. Pypaert, K. Inaba, and I. Mellman. The tetraspanin cd9 mediates lateral association of mhc class ii molecules on the dendritic cell surface. Proceedings of the National Academy of Sciences, 104(1):234–239, 2007.

40. P. A. van der Merwe. The tcr triggering puzzle. Immunity, 14(6):665–668, 2001.

41. O. S. Yousefi, M. Günther, M. Hörner, J. Chalupsky, M. Wess, S. M. Brandl, R. W. Smith, C. Fleck, T. Kunkel, M. D. Zurbriggen, et al. Optogenetic control shows that kinetic proofreading regulates the activity of the t cell receptor. eLife, 8:e42475, 2019.

## References

1. Aleksic M, et al. (2010) Dependence of t cell antigen recognition on t cell receptor-peptide mhc confinement time. Immunity 32(2):163–174.

2. Govern CC, Paczosa MK, Chakraborty AK, Huseby ES (2010) Fast on-rates allow short dwell time ligands to activate t cells. Proceedings of the National Academy of Sciences 107(19):8724–872

